# Transient delivery of A-C/EBP protein perturbs differentiation of 3T3-L1 cells and induces preadipocyte marker genes

**DOI:** 10.1101/2020.08.13.250324

**Authors:** Nishtha Sharma, Raminder Kaur, Binduma Yadav, Koushik Shah, Harshita Pandey, Diksha Choudhary, Prateek Jain, Aanchal Aggarwal, Charles Vinson, Vikas Rishi

**Affiliations:** National Agri-Food Biotechnology Institute (NABI), Knowledge City, Sector 81, Mohali, Punjab 140306, India; Department of Biotechnology, Panjab University, Chandigarh, Punjab 160014, India; Regional Centre for Biotechnology (RCB), Faridabad, Haryana 121001, India; National Cancer Institute, National Institutes of Health, MD 20892, USA

**Keywords:** C/EBPβ, Protein transfection, Adipogenesis, 3T3-L1, dedifferentiation, obesity

## Abstract

Transformation of committed 3T3-L1 preadipocytes to lipid-laden adipocytes involves timely appearance of numerous transcription factors (TFs), foremost among them C/EBPβ, is expressed during early phases of differentiation. Here we describe liposome-mediated protein transfection approach to rapidly downregulate C/EBPβ by A-C/EBP protein inhibitor. Signals from tagged A-C/EBP were observed in 3T3-L1 cells within 2hrs of protein inhibitor transfections whereas for gene transfection signals appeared in 48hrs. Following transient transfections, expression profiles of 24 marker genes belonging to pro- and anti-adipogenic, cell cycle, and preadipocytes pathways was analyzed. mRNA and protein expression profiles of adipocyte-marker genes showed lower expression in both A-C/EBP protein and gene transfected samples. Interestingly, for preadipocytes and cell fate determinant genes, striking differences were observed between protein and gene transfected samples. Preadipocyte differentiation factors Stat5a and Creb were downregulated in A-C/EBP protein samples. Five preadipocyte markers, namely, *Pdgfrα, Pdgfrβ, Ly6A, CD34 Itgb1* showed high expression in protein samples whereas only *Ly6A* and *CD34* were expressed in gene transfected samples. *Pdgfrα* and *Pdgfrβ*, two known cell fate markers were expressed in protein transfected samples 5-days post-differentiation suggesting a possible reversal of differentiation. Our study provides evidences for the robust and efficient knockdown of C/EBPβ protein to understand time-dependent gene regulation during adipogenesis.

## 1. Introduction

With an increase in severe obesity load world over and co-morbidities associated with it, there is an unprecedented interest in adipocyte biology. Adipogenesis is defined as the formation of lipid-laden adipocytes from mesenchymal stem cells. The terminal differentiation process of formation of adipocytes from committed preadipocytes is closely mimicked *ex vivo* by 3T3-L1 cell lines when induced with hormone cocktail [1]. Previous evidences have shown the pivotal role of peroxisome proliferator-activated receptor gamma (PPARγ) and members of CCAAT/enhancer binding protein (C/EBP) family in the entire terminal differentiation process [2]. C/EBPβ and C/EBPδ are first to express after induction upon hormone cocktail and are known to direct the process of differentiation by transcriptionally activating promoters of *C/EBPα* and *PPARγ*, which further lead to expression of adipocyte-specific genes and subsequent terminal differentiation [3,4]. Knocking out *C/EBPα* and *C/EBPβ* result in impaired adipogenesis whereas knocking down *C/EBPβ* by siRNA inhibited mitotic clonal expansion (MCE) which is prerequisite for adipogenesis [5,6]. Targeting members of C/EBP family of bZIP transcription factors (TFs) at initial stages of adipogenesis may bring changes in final morphology and metabolic state of adipocytes or it may inhibit differentiation process. Strategies used for inhibiting bZIPs such as use of small molecule inhibitors, siRNA, and nucleases like CRISPR/Cas, though are effective but suffer from limitations of cytotoxicity, non-specificity and off-targets effects [7– 9]. Another strategy to functionally inactivate a gene is by over-producing inhibitory variant of the same gene, known as dominant-negative. C/EBPβ activity is regulated in cells by CHOP-10, a natural dominant-negative that heterodimerize with C/EBPβ and delays its DNA-binding and invigorate preadipocytes to two cycles of MCE followed by the expression of anti-mitotic C/EBPα [10]. A-C/EBP, a designed repressive protein of C/EBP has N-terminal acidic extension appended to C/EBP leucine zipper. It forms stable heterodimer with wild type C/EBPβ, preventing its entry into nucleus thus blocking MCE and inhibiting adipogenesis [11,12]. Considering the ability of A-C/EBP to specifically inhibit C/EBPβ *in vivo*, we decided to use A-C/EBP protein and A-C/EBP plasmid to decipher the time-dependent role of C/EBPβ and its interacting partners in preadipocytes differentiation

As the process of adipogenesis involves spatial and temporal cascade of gene expression, the protocols using external DNA or RNA for transfections seem bit moderate. Usually, the genetic methods of inhibiting the expression of a protein take 24-48 hours and during the time-frame cellular and molecular requital may occur, blocking the expected phenotype [13]. Expression and temporal levels of proteins are difficult to control with nucleic acid delivery strategies since amplification steps cannot be controlled accurately [14]. Also DNA-mediated protein overexpression perturbs metabolic state of cells by promoting non-specific interactions [15]. Protein delivery is independent of transcription and translation machinery of cells and act immediately unlike gene expression from nucleic acids [16]. Protein transfection titration experiments may delineate specific interaction from non-specific interactions, can be used as a tool for basic research and may have therapeutic applications. A-C/EBP is negatively charged and highly anionic proteins or adding a polyanionic chain to proteins facilitate their delivery by same electrostatic-driven complexation as seen for nucleic acids by common cationic liposomal reagents [17].

In this study, we hypothesized that during early wave of adipogenesis if C/EBPs activity is inhibited immediately and timely during ongoing differentiation process, it may lead to changes in the original morphology and metabolic state of the cells. Inhibition of C/EBPβ may affect the expression of its interacting partners and the genes they regulate during adipogenesis. It is known that the expression of C/EBPβ takes place within few hours of induction therefore direct delivery of inhibitor can sequester C/EBPβ and bring changes in the differentiation process. Inhibition of DNA binding activity of C/EBPβ by A-C/EBP lead to lower expression of its target genes responsible for preadipocytes differentiation [11]. In this study, we used lipid-mediated transfection of negatively charged A-C/EBP protein into differentiating 3T3-L1 cells. We explored the effect of A-C/EBP on the gene expression of 24 genes involved in the process of adipogenesis [18]. Parallelly, 3T3-L1 cells were transfected with plasmid encoding A-C/EBP gene. We found that protein delivery leads to more specific and on time effects in comparison to genetic methods as later take hours to days for translating the protein. Initial gene expression analysis of common preadipocyte marker genes namely, Platelet-derived growth factor receptor α (*Pdgfrα*), Platelet-derived growth factor receptor β (*Pdgfrβ*), Lymphocyte antigen six complex, locus A (*Ly6A*), *CD34* and fibronectin receptor beta subunit (*Itgb1*) have shown increased expression only in protein transfected cells whereas A-C/EBP gene transfected cells showed upregulation of *Ly6A*, and *CD34*. Direct delivery of purified A-C/EBP protein inside committed preadipocytes has led to their non-conversion into adipocytes even after hormonal induction. Gene expression analysis suggest change in the cell fate and reversal of differentiation of 3T3-L1 cells when exposed to A-C/EBP protein during early phase of differentiation.

## 2. Materials and Methods

### 2.1 Cell culture and differentiation

3T3-L1 cell lines were obtained from NCCS, Pune (India) at passage number 16. Cells were were cultured and maintained in DMEM supplemented with 10% (vol/vol) bovine serum, 100U/ml penicillin and 100μg/ml streptomycin, in 5% CO_2_ incubator with 98% humidity. Cells were grown till 60-70% confluency before differentiation and transfection experiments. Cells were differentiated on day 0 with differentiation media containing DMEM supplemented with 10% FBS and MDI (0.5mM 3-iso-butyl-1-methylxanthine, 1μM dexamethasone and 1μg/ml insulin) for 48hrs. After 48hrs, differentiation media was replaced with adipocyte maintenance media containing DMEM supplemented with 10% FBS and 1μg/ml insulin. After day 4, cells were fed every other day with DMEM supplemented with 10% FBS. Cells were seeded in 60mm dishes for transfection and differentiation experiments. Transfected cells were collected post differentiation for RNA and protein isolation and were subjected to mRNA and Western blotting experiments.

### 2.2 Cloning, protein expression and purification

Cloning of dominant-negative A-ZIPs i.e. A-C/EBP and A-VBP in bacterial expression system has been described earlier [19]. A-ZIPs (A-C/EBP and A-VBP) were cloned as KpnI-EcoRI fragments into pCMV, a mammalian vector [20]. For localization studies, EGFP was cloned as HindIII-NotI fragment immediately downstream of A-C/EBP in pT5 vector. Vectors containing dominant-negative genes were used to transform *Escherichia coli* (BL21 DE3), grown overnight at 37°C in Super Broth containing 100 μg/ml ampicillin and 35μg/ml chloramphenicol. 20ml primary cultures were transferred to 500 ml Super Broth containing 100 μg/ml ampicillin and grown until OD reached 0.6. Cultures were then induced with 1mM isopropyl-β-D-thiogalactopyranoside (IPTG) and harvested after 3hrs and processed as described earlier [21]. Dialysis was performed against low salt buffer (20mM Tris-HCl pH 8.0, 50mM KCl, 1mM EDTA, 0.2mM phenylmethylsulfonyl fluoride (PMSF), and 1mM Dithiothreitol (DTT). Dialyzed A-C/EBP and A-VBP proteins were passed through hydroxyapatite column and eluted with dialysis buffer containing 250 mM sodium phosphate buffer (pH 7.4). Proteins were further purified to high homogeneity using reverse phase analytical HPLC system equipped with C-18 hydrophobic column as described earlier [21]. Lyophilized HPLC purified proteins were dissolved in phosphate buffer saline (PBS) with EDTA and resolved on SDS-PAGE to check the purity and quality (Fig S1A). Proteins were quantified using UV spectroscopy and concentrations were calculated according to previously published method [21]. HPLC purified proteins were used for mammalian cell line transfection experiments. A-C/EBP-EGFP was also purified following same protocol and quality was checked with SDS-PAGE after passing through hydroxyapatite column. After dialysis protein was used for transfections (Fig S1B).

### 2.3 Liposome-mediated cell transfection and confocal microscopy

Two days prior transfections, the cells were seeded in 6-well plates/60mm dishes for RNA isolation and immunoblotting, respectively. For plasmid transfections, 2.5μg of vector was added to 50μl low serum media (LSM), followed by addition of 5μl of Lipofectamine 2000 in 50μl LSM. The plasmid and Lipofectamine mixture was incubated for 15-20 minutes at room temperature (RT) and then added to the cells. MTT [3-(4,5-<u>dimethylthiazol</u>-2-yl)-2,5-diphenyltetrazolium bromide] assays were performed to check cell-viability at different concentrations (0.1 μM-20 μM) of pure proteins. For A-C/EBP protein transfections, pure protein was added to 50μl LSM so that the final concentration was 3μM, followed by addition of 50 μl LSM containing 5 μl of Lipofectamine 2000. After incubation for 20 minutes at RT, the mixture was added to the cells. The work-flow for protein and gene transfections and differentiation are given in supplementary section (Fig S2A). Similarly, A-C/EBP-EGFP protein was used to transfect the cells on culture slide and after two hours, cells were fixed and treated with Hoechst stain. Purified A-C/EBP (30 - 40 μg) was tagged with Alexa Fluor 488 according to manufacturer’s protocol (Thermo fisher). Time lag expression of A-C/EBP protein was observed using EGFP tagged A-C/EBP plasmid (pCMV-A-C/EBP-EGFP) to transfect the cells on culture slide (2hrs and 48hrs). Fixed cells were visualized and photographed using confocal microscopy (Carl-Zeiss LSM 880).

### 2.4 Oil Red O staining

For Oil Red O (ORO) staining, 3T3-L1 cells were washed three times with PBS and fixed for 20 min using 3.7% formaldehyde. Oil red O stain (0.5% in isopropanol) was diluted with water (3:2) and filtered through 0.45μm filter. Cells were incubated with filtered ORO for 1hr at RT. Stained cells were washed 2-3 times with water and lipid droplets were visualized by light microscopy and were photographed. ORO stained cells were eluted in 100% isopropanol and signals quantified by taking absorbance at 520nm. Neat isopropanol was used as blank.

### 2.5 Protein estimation

The protein contents were estimated by bicinchonic acid (BCA) method using the Puregene BCA protein assay kit.

### 2.6 Immunoblotting

Cell monolayers (6 cm dishes), at different time points were washed with cold PBS, pH 7.4 and then scraped in presence of RIPA buffer (150mM NaCl, 0.1% Triton X-100, 0.5% Sodium deoxycholate, 0.1% SDS, 50mM Tris Cl pH 8.0 and protease inhibitors). Cell lysates were cleared by centrifugation and supernatants were heated at 98°C for 10-12 minutes. Equal amounts of protein were separated by 15% SDS/PAGE along with pre-stained protein ladder. Gel-resolved proteins were transferred to PVDF membranes in cool conditions for 2 hours in transfer buffer (25mM Tris, 190mM glycine, 20% methanol, pH8.3). Membrane was blocked using 3% BSA in TBS containing 0.1% Tween-20 for 1hr at RT. T7-tag HRP-conjugated antibody (Thermo fisher, Cat. no. PA1-31449) was used at a dilution of 1:10000 for overnight at 4°C and probed for protein using ECL chemiluminescence kit (BioRad). HPLC purified protein was taken as control. Parallel blot with same amount of protein was run and probed for β-actin under same conditions (Cell Signalling Technologies, Cat. no. 4967S). β-actin band was used as a loading control. Immunoblotting was performed for C/EBPβ expression in cells transfected with plasmid containing A-C/EBP gene and pure A-C/EBP protein. Anti-C/EBPβ antibody (Biolegend, Cat. no. 606202) was used at 1:2000 dilution for overnight at 4°C. Blots were washed with TBST and incubated with anti-mouse (Cell signalling technologies, Cat. no. 7076P2) horse radish peroxidase conjugated secondary antibody for 2hrs at RT. For C/EBPδ, anti-C/EBPδ antibody (Cloud clone, Cat. no. PAC462Mu01) was used overnight at 1:1000 dilution and 4°C. Parallel blot with same amount of protein was run and probed for β-actin under same conditions. For adipocyte-marker genes, protein was isolated from transfected cells after four days of differentiation. Anti-C/EBPα (Cell signalling technologies, Cat. no. 8178S), anti-PPARγ (Cloud clone, Cat. no. PAA886Mu01), and anti-aP2 (Cloud clone, Cat. no. PAB693Mu01)) were used at a dilution of 1:2000, 1:1000, 1:1000 respectively, for overnight incubation. Blots were incubated with anti-rabbit (Cell signalling technologies, Cat. no. 7074P2) secondary antibody for 2hrs at RT. Parallel blot with equal amount of protein sample was used to probe with anti-β-actin antibody as loading control under same conditions. Proteins were visualized by ECL chemiluminescence method using Amersham imager 600 (GE healthcare) and densitometry of protein bands was performed with ImageJ software.

### 2.7 RNA isolation, qRT PCR and protein-protein interaction network prediction

For gene expression study, transfected cells were lysed with TriZol reagent and total RNA was isolated as per manufacturer’s instructions. Isolated total RNA was treated with DNaseI to remove traces of genomic DNA. 1μg of total RNA was reverse transcribed to cDNA in a final reaction volume of 20μl using BioRad iScript cDNA synthesis kit. qRT PCR was performed with gene-specific primers (Table S1) using iTaq universal SYBR Green Supermix (BioRad) for 45 cycles on CFX96 Real time system (BioRad). The C_t_ values obtained were normalized against actin because of its consistent expression as observed in Western blots. qRT-PCR data was analyzed by comparative C_t_ method [22]. The factors involved in adipogenesis and mitotic clonal expansion were selected from literature[11,18]. Selected factors were put in STRING database in Multiple protein sections by names/identifiers section and interaction data was retrieved. For preadipocyte-marker genes, RNA was isolated from transfected cells 5-days post-differentiation and qRT-PCR was performed using gene specific primers retrieved from Harvard Primer Bank (Table S3).

### 2.8 Chromatin immunoprecipitation analysis

For analysing the binding of C/EBPβ to the promoters of adipogenic genes, ChIP assay was performed as described earlier with minor modifications [23]. Briefly, cells transfected with gene and protein along with untreated cells were cross-linked with 1% formaldehyde for 15min at RT. The cross-linking reaction was stopped with addition of 0.125M glycine for 5min. Cells were washed with ice-cold PBS containing protease-inhibitor cocktail (PIC) and centrifuged at 3000 rpm for 5min. Pellet was suspended in lysis buffer (50mM HEPES-KOH (pH-7.5), 140mM NaCl, 1mM EDTA pH-8, 1% TritonX-100, 0.1% sodium deoxycholate, 0.1% SDS, protease inhibitor cocktail) and kept on ice for 10 min. Suspended cells were sonicated for 8 cycles (30s on 30s off) with 30% amplitude on ice using probe sonicator (Sonics vibra cells, Fisher scientific) followed by centrifugation at 9000 rpm for 10 min. RNA and protein-free DNA samples were obtained by adding RNase and proteinase K to 50μl of sample at 65°C for 2 hours and purifying the DNA using PCR purification kit (Thermo fisher scientific). Purified DNA was run on 1.5% agarose gel to check efficiency of chromatin shearing. 2μg of chromatin was used for immunoprecipitation (IP). Samples were incubated at 4°C for overnight with anti-C/EBPβ and non-specific Rabbit IgG. Sample without IP were used was used as input control. Next day, Protein G beads (G Biosciences) were vortexed and added to Antibody-chromatin samples for 2hrs at 4°C with rotation. Beads were pelleted down using Magnetic separation rack (NEB,USA) and washed with low salt buffer followed by high salt buffer and LiCl buffer for 5min each at 4°C. Elution buffer (1% SDS, 100mM NaHCO_3_) was added to the beads and input samples and incubated for 30min at 65°C. Eluted chromatin was reverse cross linked by adding 5M NaCl and Proteinase K for 2hrs at 65°C. Samples were passed through spin column and eluted DNA was used for semi-quantitative or qRT-PCR for 32-40 cycles with specific primers (Primer sequences are given in Table S2). ChIP amplified 150-200 bp DNA fragments containing C/EBP-binding site present in the promoter regions of adipocyte marker genes [24]. Analysis of quantitative PCR results was done by using fold enrichment method [(Fold enrichment= 2^-ddCt^) where ddCt= Ct (IP)-Ct (IgG)].

### 2.9 Statistical analysis

For gene expression analysis, means and standard errors were calculated from six biological replicates. The statistical significance was determined by Student’s *t* test. For Western blotting, means and standard errors were calculated from three biological replicates. The statistical significance was determined by one-way ANOVA. A difference in p value of <0.05 is considered as significant.

## 3. Results

### 3.1 Cellular uptake of A-C/EBP protein and its impact on lipid accumulation of murine 3T3-L1 cells

Before embarking on performing differentiation experiments, cytotoxic effect of A-C/EBP protein was tested on 3T3-L1 cells by performing MTT assay in concentrations range 0.1-20μM (Fig S3A). Percentage survival at 3μM was 80%, confirming the non-toxicity of A-C/EBP. Three experimental conditions were used to observe the time-dependent appearance of fluorescent tagged A-C/EBP protein signals in 3T3-L1 cells. 1) Cells transfected with A-C/EBP protein for 2hrs. 2) A-C/EBP plasmid transfected for 2hrs followed by washing and replenishing the cells with fresh media without plasmid. 3) 3T3-L1 cells were transfected with A-C/EBP plasmid for 48 hrs. Confocal microscopy was used to demonstrate the cellular uptake and localization of A-C/EBP in 3T3-L1 cells. Both Alexa Fluor 488 (Al Fl 488) tagged A-C/EBP and A-C/EBP-EGFP (37kDa) proteins were able to internalize into the cells and intense A/CEBP protein signals were observed within 2 hrs. (Fig 1A). Arrows depict the presence of tagged A-C/EBP in the cytoplasm of 3T3-L1 cells. Cells transfected with pCMV-A-C/EBP-EGFP plasmid for 2hrs and 48hrs showed very low expression of EGFP in 2hrs that intensified in 48hrs and became equivalent to 2hrs protein samples (Fig S2). Our results show that liposomes can carry and deliver large anionic protein A-C/EBP-EGFP (37kDa) inside the cells. Furthermore, we determined the impact of A-C/EBP on the morphology of 3T3-L1 cells after differentiation. 60-70% confluent preadipocytes were treated with increasing concentrations of A-/CEBP ranging from 0.1-9μM. 10 days, post-differentiation, cells showed lower lipid accumulation with increasing concentration of A-C/EBP as evidenced by ORO-staining compared to untransfected cells (Fig1B). Quantification of ORO stained cells showed a linear decrease in lipid accumulation with increasing concentrations of A-C/EBP (Fig 1C). In another set of experiments, preadipocytes treatment with A-C/EBP protein (3μM) for 2hrs and A-C/EBP plasmid (1μg) for 48hrs led to equivalent decrease in lipid accumulation (Fig 1D-E) suggesting that plasmid needed approximately 48hrs to transcribe and translate A-C/EBP protein. Preadiocytes transfected with A-C/EBP plasmid achieved similar reduced differentiation grade as protein transfections but only after a lag of two days. Transfection with A-VBP, a dominant negative inhibitor of VBP, which is not reported to play a role in the process of adipogenesis, showed no change in lipid accumulation compared to control cells (Fig S2B, [12]).

**Figure 1:**
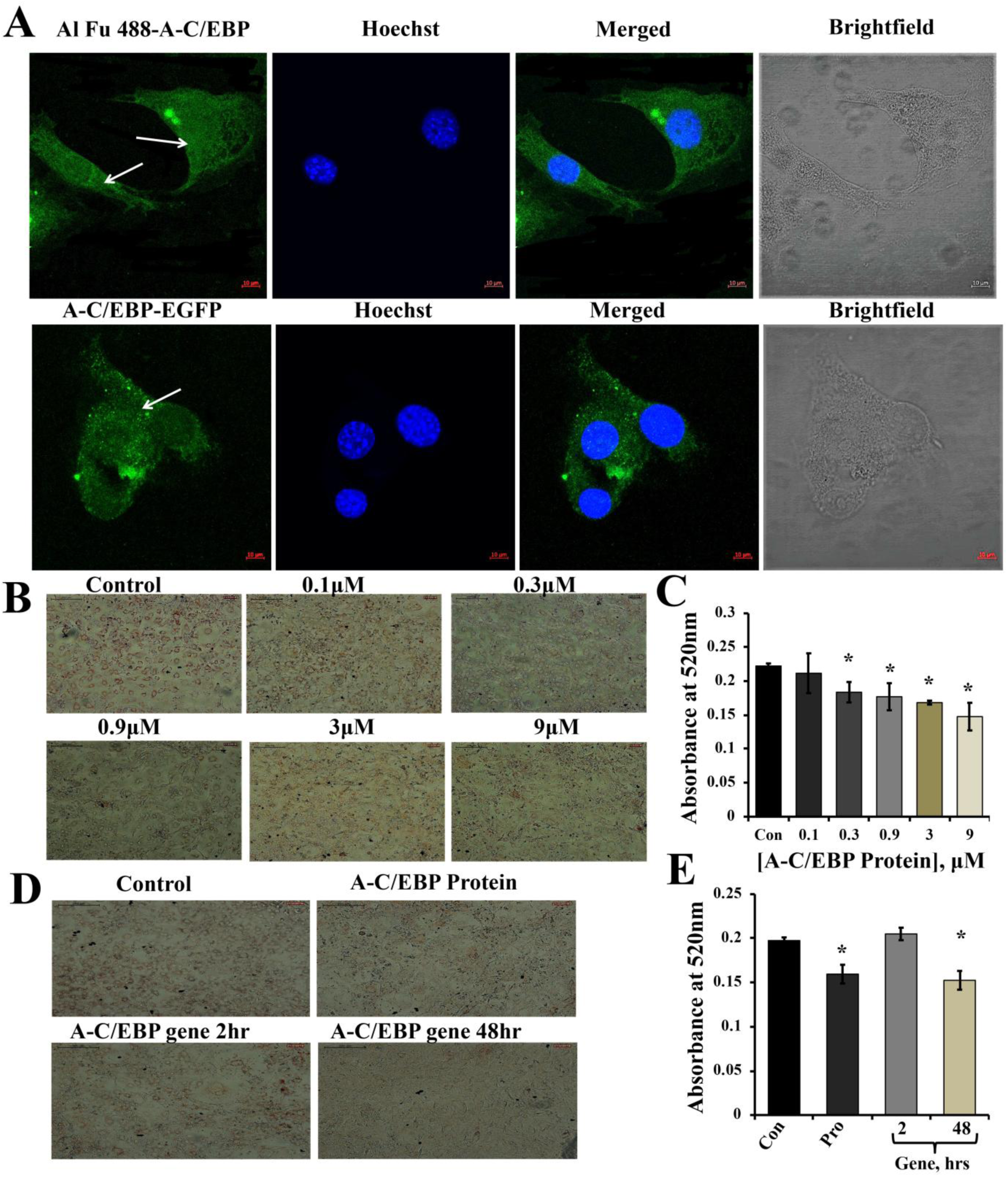
Localization of protein in cytoplasm and ORO staining of transfected and differentiated cells. (**A**) Image representing Alexa fluor (Al Fu) 488 and EGFP tagged protein inside the cytoplasm, arrows depict the presence of tagged A-C/EBP in the cytoplasm, counterstained with Hoechst 33342; Magnification =40X, Scale bar =10μm. (**B**) Representation of ORO stained cells transfected with A-C/EBP protein along with untreated cells. Half-log concentrations of A-C/EBP protein were taken. Magnification =10X, Scale bar =200μm. (**C**) Quantification of ORO stained samples treated with A-C/EBP protein. Absorbance was taken at 520nm, and isopropanol was taken as blank. (**D**) Image representing ORO staining of cells transfected with different concentrations of A-C/EBP protein and gene; Magnification=10X, Scale bar =200μm. (**E**) Quantification of ORO stained samples treated with A-C/EBP protein and gene, absorbance was taken at 520nm, isopropanol was taken as blank. *Significantly different from control (p<0.05).

### 3.2 Time-dependent appearance of A-C/EBP protein in the cells and lower expression of adipocyte-specific markers in instilled protein samples

3T3-L1 cells were transfected with A-C/EBP protein and gene for different time intervals, as indicated in figure 2A-B. Western blots analysis showed A-C/EBP gene expression only after 24hrs, peaked at 48hrs and persisted till 72hrs (Fig 2B), whereas, the levels of purified A-C/EBP protein delivered directly into the cells, peaked in just 2hrs of transfection, then decreased rapidly and was barely detectable after 10hrs (Fig 2A). The trend lines showed the increasing and decreasing expression of A-C/EBP in protein and gene transfected cells at different time intervals (Fig 2, C-D). Prompted by immunoblotting experiments, cells were transfected with A-C/EBP protein for 2hr, 4hr and 6hr and then differentiated with MDI hormone cocktail. RNA was isolated from these samples and qRT-PCR was performed for *C/EBPβ, C/EBPδ* and adipocyte-marker genes (*C/EBPα, PPARγ, 422/ap2, GLUT4, SREBP1, Scd1*). Results showed no significant change in gene expression of *C/EBPβ* and *C/EBPδ* within 2hr and 4hr of transfected and differentiated cells (Fig 2E). At 6hr, lower expression of *C/EBPβ* was observed. C/EBPβ autoregulates its own transcription by binding to its promoter region located between −121 and −71 [25]. We observed that C/EBPβ mRNA expression was not affected by A-C/EBP protein in 4hrs but in 6hrs *C/EBPβ* expression decreased which suggests degradation of C/EBPβ|A-C/EBP heterodimer leading to hampering of autoregulation of transcription. The mRNA expressions of six adipocytes-marker genes i.e., C/EBPα, PPARγ, 422/ap2, Glut4, SREBP1, and Scd1 were lower in all three experimental conditions of A-C/EBP protein transfection (Fig 2F-H).

**Figure 2:**
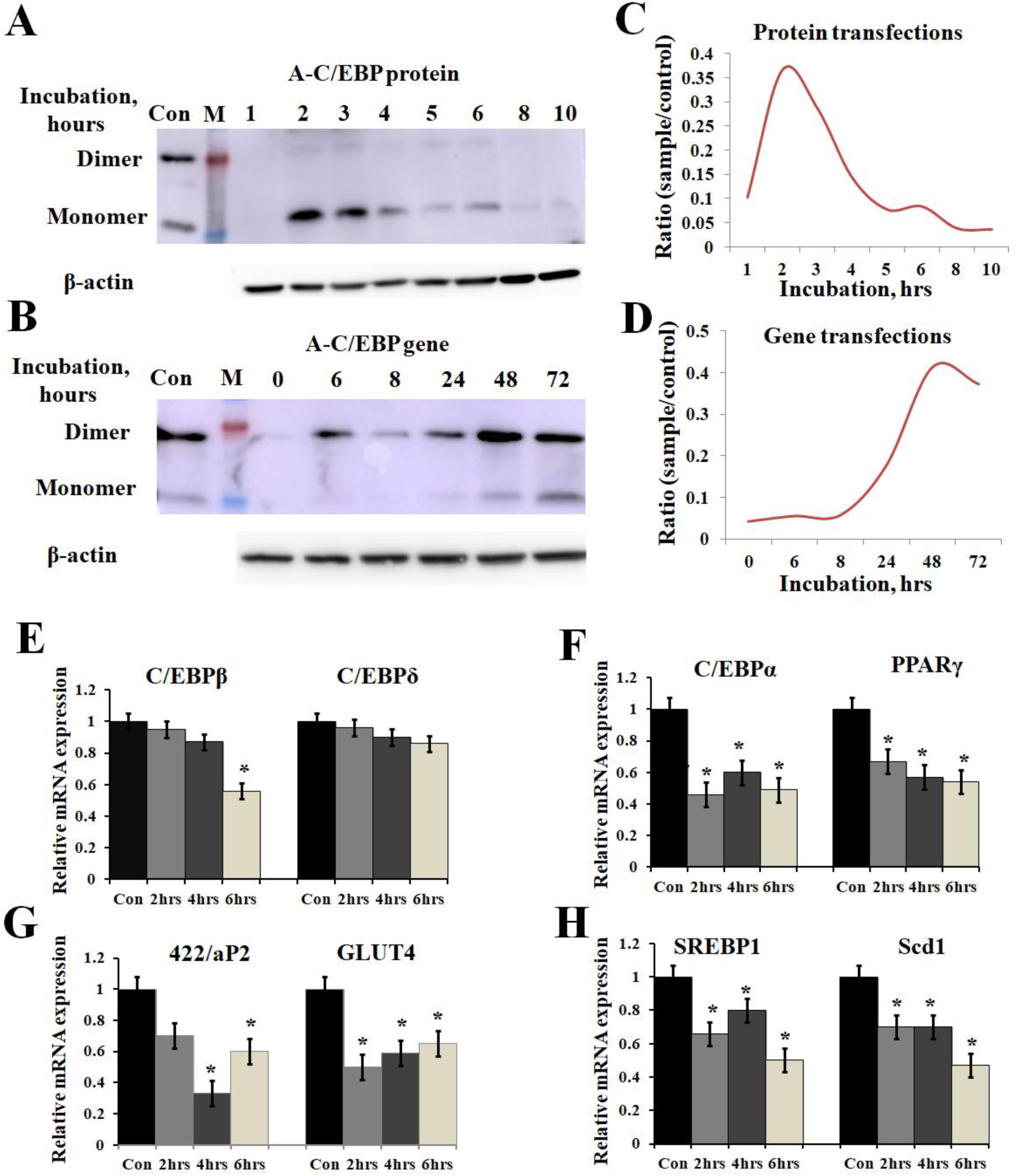
Time-dependent presence of A-C/EBP protein and A-C/EBP gene product in 3T3-L1 cells and effect of A-C/EBP protein on adipocyte marker genes. (**A**) Western blots show the signals of A-C/EBP protein when the cells were treated with liposome entrapped A-C/EBP protein. With time, signal intensity decreased due to degradation of A-C/EBP protein. Pure protein was used as control. (**B**) Western blot shows the time-dependent profile of A-C/EBP gene product. Here the signal intensity increased and peaked at 48hrs and remained so at 72 hrs. Pure protein was used as control. (**C-D**) Curves show A-C/EBP signal trends plotted from Western blot experiments over indicated time. Opposite trends were obtained in protein and gene transfected cells. *Significantly different from control group (p < 0.05). Values are expressed as mean ±SD; n=3 independent experiments. (**E-H**) Real time PCR analysis of C/EBPs and *PPARγ, 422/aP2, GLUT4, SREBP1, Scd1* w.r.t actin in cells transfected with A-C/EBP for 2hr, 4hr and 6hr before differentiation. *Significantly different from control (p<0.05). Values are expressed as mean ±SD; n=6 independent experiments.

Primary embryonic fibroblasts from double knockout *C/EBPβ*(-/-).*δ* (-/-) mice, neither differentiate nor express the late TFs i.e., C/EBPα and PPARγ [6]. Knocking down *C/EBPβ* with A-C/EBP expressing gene has resulted in impaired adipogenesis, prevents MCE and inhibit the expression of adipocyte-marker gene *422/aP2* along with *C/EBPα* and *PPARγ* [11]. We also observed lower gene expression of selected adipocyte-marker genes in A-C/EBP protein as well as gene (48 hrs) transfected cells, supporting the above observations. No significant difference in the expressions of these genes were observed in samples that were transfected with A-C/EBP gene for 6hrs. mRNA expression analysis of adipocyte-marker genes from cells transfected with negative-control A-VBP protein and gene showed no significant change in expression compared to untreated samples under three experimental conditions used here (Fig 3A-F).

**Figure 3:**
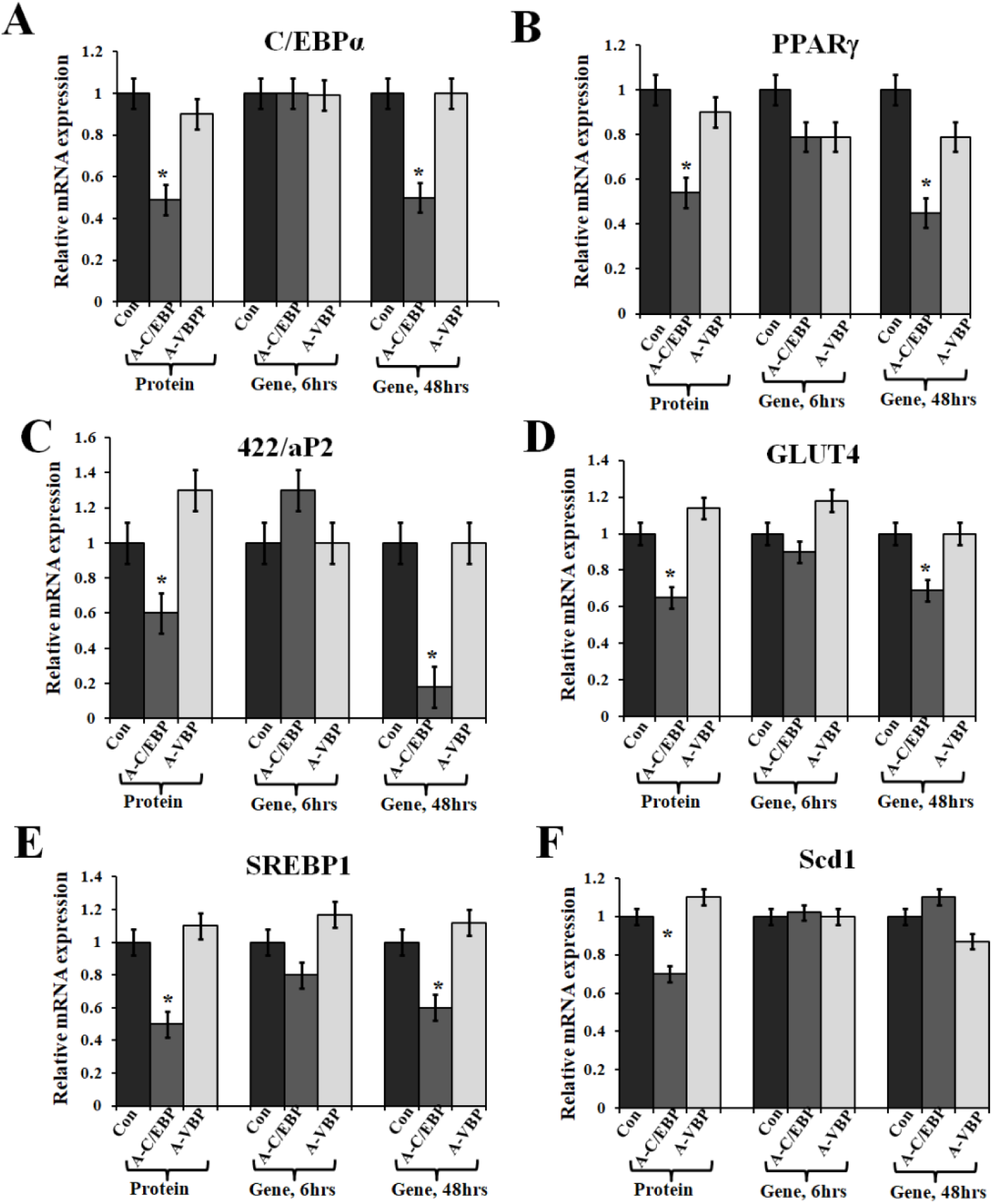
Effect of A-C/EBP and A-VBP protein and gene on six adipocyte marker genes. (**A-F**) Gene expression analysis for comparative expression of *C/EBPα, PPARγ, 422/aP2, GLUT4, SREBP1, Scd1* w.r.t. actin expression. 3T3-L1 cells were transfected with A-C/EBP and A-VBP protein. A-VBP is a designed dominant negative of VBP transcription factor and used as a negative control here. All six marker genes show downregulation in cell samples transfected with 3 μM A-C/EBP protein whereas only 48 hrs gene samples showed significant decrease. Since neither A-VBP protein nor its gene has any effect on expression of six adipocytes marker genes we conclude that A-C/EBP is specific in inhibiting preadipocyte 3T3-L1 cells differentiation. *Indicate significantly different from control (p<0.05). Values are expressed as mean ±SD; n= 6 independent experiments.

### 3.3 Alterations in adipogenic-specific TFs and preadipocyte-marker genes in transfected and differentiated 3T3-L1 cells

Taking cues from previous studies, we chose foremost transcription factors involved in adipocyte differentiation [18]. The TFs chosen for this study for gene expression analysis are both promoter and inhibitors of adipogenesis. *KLF6, Stat5a, Zfp423, Zfp467, TCF7L1, CREB* and *VEGF* are promoters of adipogenesis [26–29]. On the other hand, *Pref1(Dlk1), KLF2, GATA2/3* and *FoxC2* are known to inhibit the process of differentiation [26]. Among all the selected factors of early wave of adipogenesis, *Stat5a, CREB, GATA2/3, FoxC2, KLF2, VEGF* and *Cdk* are known to interact with *C/EBPβ* directly or indirectly [30–35]. STRING database was used to predict the putative interacting partners of C/EBPβ among these chosen TFs. Ten TFs were found to interact with C/EBPβ (Figure 4A). Five TFs showed no interaction with C/EBPβ as predicted by STRING database. To quantify the effect of A-C/EBP protein and gene on expression of major TFs involved in 3T3-L1 differentiation, cells were treated with both purified A-C/EBP protein and *A-C/EBP* coding plasmid. A-VBP was used as negative control. mRNA samples from A-C/EBP untreated and treated cells were generated after 48hrs of differentiation. Total of 24 adipogenic-specific genes and other TFs having roles in early adipogenesis (within 48hrs of differentiation) were selected and subjected to qRT-PCR analysis. It was found that expression of *C/EBPβ* was significantly lower in cells transfected with A-C/EBP protein 6hr post-transfection and *A-C/EBP* gene 48hr post-transfection compared to control (without transfection) and 6hr post-transfection gene samples. Change in *C/EBPδ* expression was not significant. Results indicate high specificity of action of A-C/EBP in inhibiting C/EBPβ. Among pro-adipogenic genes, expressions of *KLF6, Stat5a, ZFp467*, and *CREB* were lower in protein transfected cells with no change in A-C/EBP gene transfected cells (48hr). Among anti-adipogenic proteins, only *GATA2* showed higher expression in protein transfected cells whereas in gene transfected cells (48hr) the expressions of all anti-adipogenic genes except *GATA2* were higher. The expressions of selected cell cycle markers i.e., *cyclin A, cyclin B1, cyclin D1* were lower in A-C/EBP protein and gene 48hr-post transfection samples and expression of *p27* was higher than control (Figure 4B). The degradation of p27 is necessary for cell cycle to progress [11]. The high expression of *p27* and low expression of cyclins suggest the inability of preadipocytes to undergo MCE even after induction. Inhibition of differentiation may be due prevention of C/EBPβ to localize into nucleus after it heterodimerized with A-C/EBP [11,36]. In gene 6hr transfected cells, the expression of *KLF6, Stat5a, Zfp423, CREB, KLF2, GATA2, GATA3, FoxC2, VEGF* remain unchanged but the expressions of *Pref1* and *p27* were lower with higher expression of cell cycle genes. A-VBP transfected cells showed no significant changes in mRNA expressions for all genes studied here under four experimental conditions (control, protein transfected, gene 6hr post transfection and gene 48hr post transfection) (Figure 4B).

**Figure 4:**
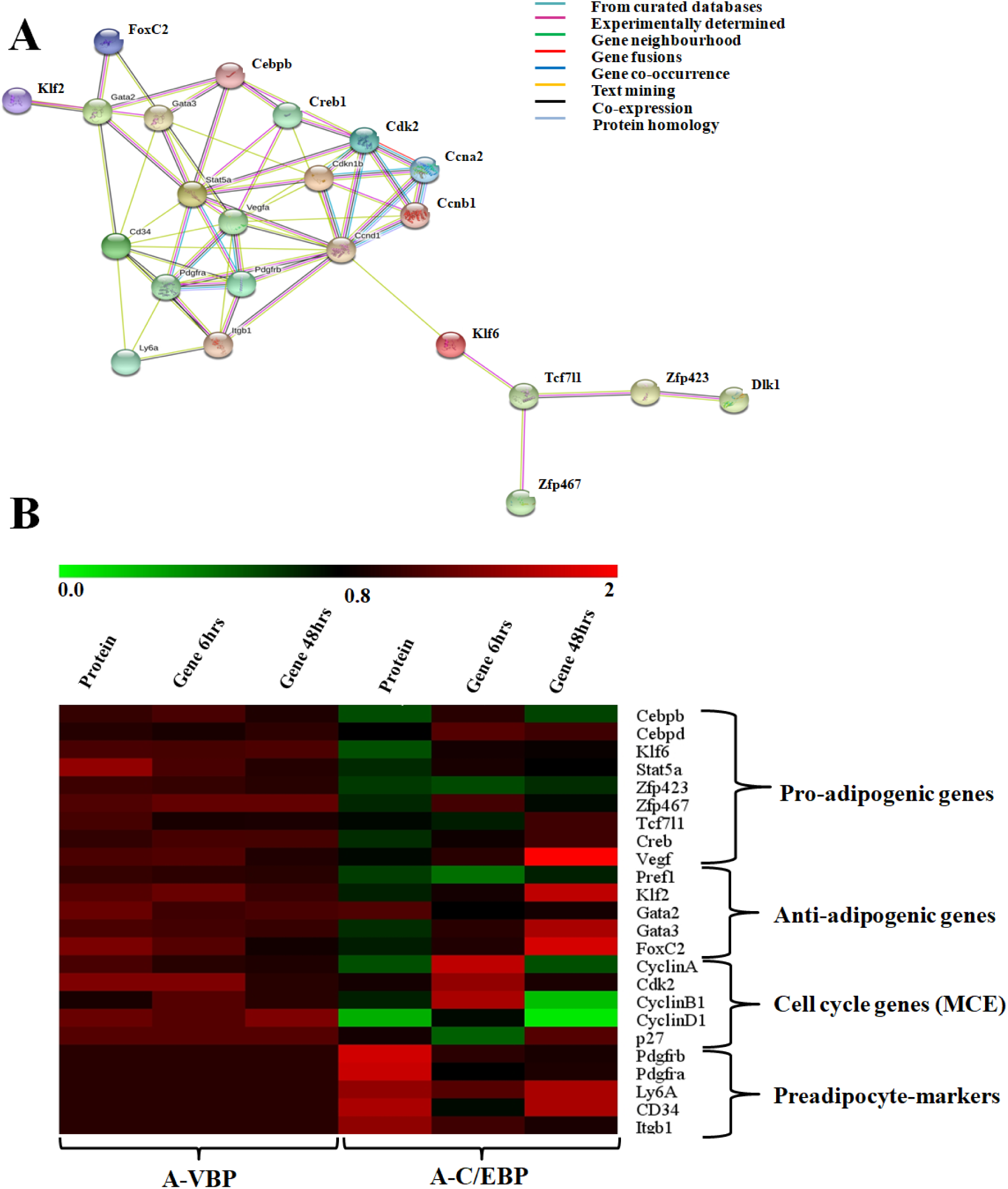
Effects of A-C/EBP and A-VBP proteins and gene transfections at the indicated conditions on the expression of factors involved in adipogenesis, MCE and preadipocyte state of 3T3-L1 cells. (**A**) Depiction of interaction of adipogenic genes with C/EBPβ using STRING database (**B**) Heat map representing Real time PCR analyses of the expression of 24 genes involved in the process of differentiation, mitotic clonal expansion and preadipocyte markers from the cells transfected with A-C/EBP and A-VBP protein and gene. The color indicates the fold-change value converted to log2 scale, as compared to untreated sample. The data represent the mean of 6 independent biological experiments and transcript were normalized to actin.

Based on the morphology of ORO stained cells and lower lipid-accumulation in A-C/EBP protein and gene (48hr) transfected cells, we performed qRT-PCR for common preadipocyte-marker genes [37–39]. Among these markers, *Ly6A, CD34* and *CD29* are common mesenchymal stem cell markers [37]. Signalling balance between *Pdgfrα/Pdgfrβ* modulate the progenitor cells to commit to either white (*Pdgfrβ*) or beige (*Pdgfrα*) adipocytes with very different metabolic profiles [40]. Analysis of qRT-PCR results showed increase in expression of all five common preadipocyte-marker genes i.e., *Pdgfrβ, Pdgfrα, Ly6A, CD34* and *Itgb1* in A-C/EBP protein transfected cells (Figure 4B). Surprisingly, in 48hr A-C/EBP gene transfected cells, though expression of marker genes *Pdgfrβ, Pdgfrα*, and *Itgb1*was not significantly different from control but *Ly6a* and *CD34* were found to be upregulated (Figure 4B). CD34, of all five selected markers, expressed in both preadipocytes and adipocytes whereas others expressed only in committed preadipocytes [38].

### 3.4 Effect of A-C/EBP on protein expression of C/EBPs and adipocyte markers

A-C/EBP protein was designed to inhibit the DNA binding of all C/EBP family members [12]. A transgenic mice overexpressing A-C/EBP protein showed reduced C/EBPβ expression but elevated C/EBPα level that suggest higher specificity of action in *in vivo* model [41]. Earlier it was shown that the heterodimer between peptide inhibitor A-ZIP and its targeted bZIP (bZIP|A-ZIP) is degraded by the proteolytic machinery of the cell [41,42]. To determine the effect of A-C/EBP on expression of C/EBPβ, total protein was isolated from gene and protein transfected cells. The relative intensity of bands was estimated with respect to β-actin (Fig 5B-C). C/EBPβ expression (C/EBPβ and LAP isoforms, 35 and 32kDa) was found to be 36% lower in 2hrs A-C/EBP protein samples, and around 38% lower in 48hrs A-C/EBP gene samples (Fig 5A). In contrast, LIP form (20kDa) was found to be upregulated in gene transfected cells in 6hrs and 48hrs samples. The protein expression of C/EBPδ was not affected by neither A-C/EBP gene or protein transfections (Figure 5A-B). The third member of C/EBP family i.e., C/EBPα (42kDa) was downregulated in protein and 48hr gene transfected samples, although the expression was much lower in gene transfected compared to protein samples (Figure 5C). Similarly, expression of 422/aP2 also follows similar trend as that of C/EBPα (Figure 5C). For PPARγ, the decrease in expression in both protein and 48 hr gene transfection was comparable (Figure 5C). Degradation of C/EBPβ in early stages of differentiation led to lower expression of adipocyte-marker genes. Major adipocyte-marker genes such as *PPARγ, C/EBPα* and *422/aP2* exhibited lower protein expression (Figure 3A-C, 5C-D). As loading control, the blot membrane was incubated with anti-β actin antibody.

**Figure 5:**
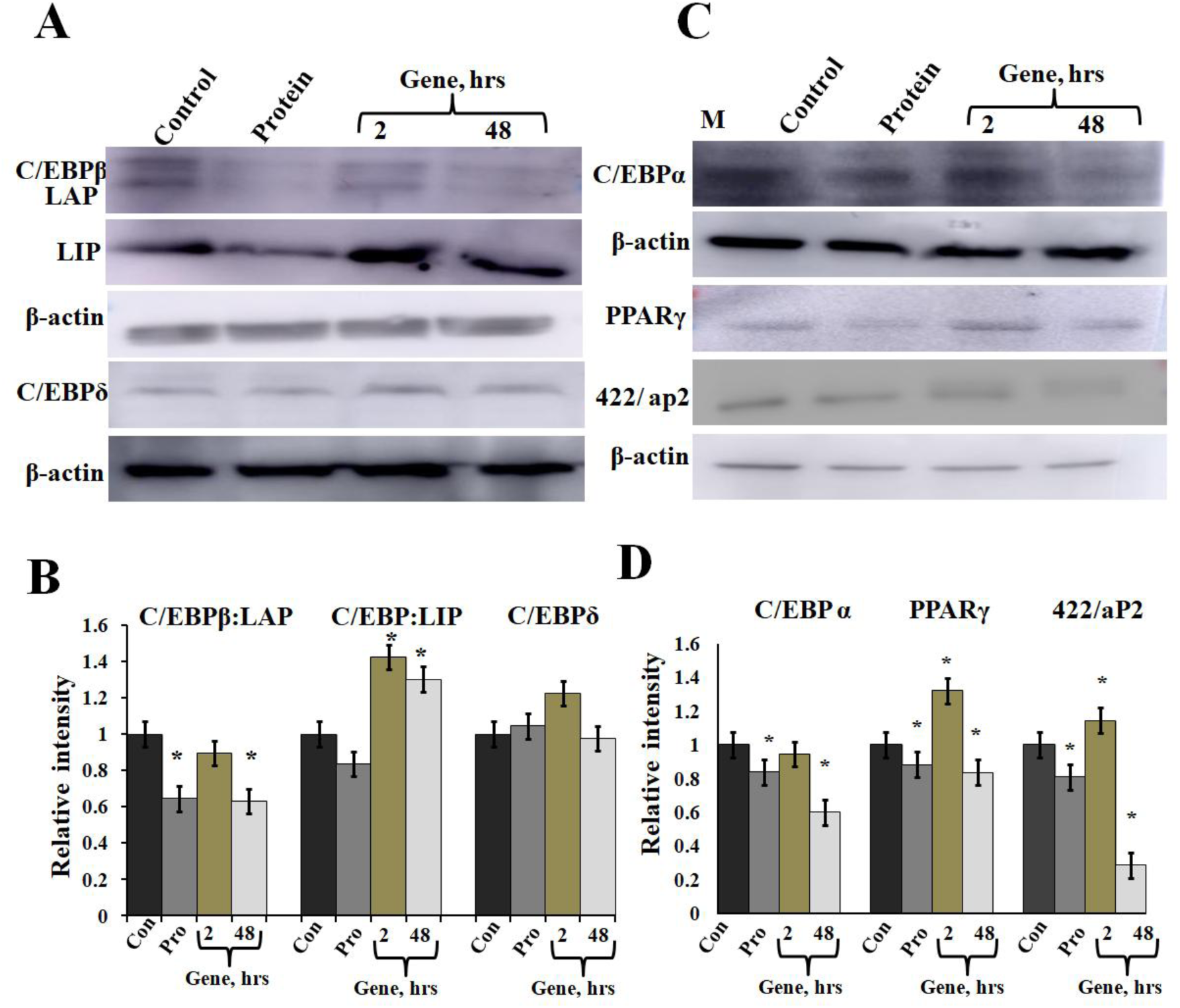
Expression of C/EBPβ, C/EBPδ and adipocyte marker genes in cells transfected with A-C/EBP protein and gene. (**A**) Immunoblotting of C/EBPβ (C/EBPβ 35kDa, LAP 32kDa and LIP 20kDa) and C/EBPδ (30kDa) from protein isolated from A-C/EBP protein and gene transfected cells. (**B**) Relative quantitative analysis of C/EBPβ, LAP, LIP isoforms and C/EBPδ w.r.t actin gene. (**C**) Immunoblotting of adipocyte-marker genes C/EBPα (42kDa), PPARγ (53 and 57kDa), 422/aP2 (16kDa) from protein and gene transfected cells. (**D**) Relative quantitative analysis of C/EBPα, PPARγ, 422/aP2 w.r.t actin. *Significantly different from control group (p < 0.05). Values are expressed as mean ±SD; n=3 independent experiments

### 3.5 A-C/EBP treatment of cells led to reduced binding of transcription factor at C/EBP regulatory elements in adipocyte-marker genes except C/EBPα

Chromatin immunoprecipitation followed by semi-quantitative and qRT-PCR was performed for transcription factor binding at promoter site of adipocyte-marker genes. Earlier studies showed that C/EBPβ act as transcriptional activator of both *C/EBPα* and *PPARγ* [4,36]. After acquiring DNA-binding activity, C/EBPβ binds to regulatory elements in promoters of *C/EBPα, PPARγ* and *422/aP2* genes [24]. To determine C/EBPβ binding to promoter region of these genes, ChIP assay was conducted using cells transfected with A-C/EBP gene and protein. Cells were cross-linked and immunoprecipitated with anti-C/EBPβ antibody and PCR primers corresponding to adipocyte-marker genes were used. Following A-C/EBP gene and protein transfections, reduced binding was observed for C/EBPβ at promoter-region of *PPARγ* and 422/aP2 in comparison to untransfected control and 2hrs gene transfections (Figure 6A). Interestingly, the amplification of promoter-region of C/EBPα was enhanced in both protein and gene transfections. We construe this observation as due to enhanced binding of C/EBPα to its own promoter-region, although enhanced binding did not lead to increased protein expression [24]. The fold enrichment w.r.t. non-specific antibody agrees with the results of semi-quantitative PCR (Figure 6 B-D).

**Figure 6:**
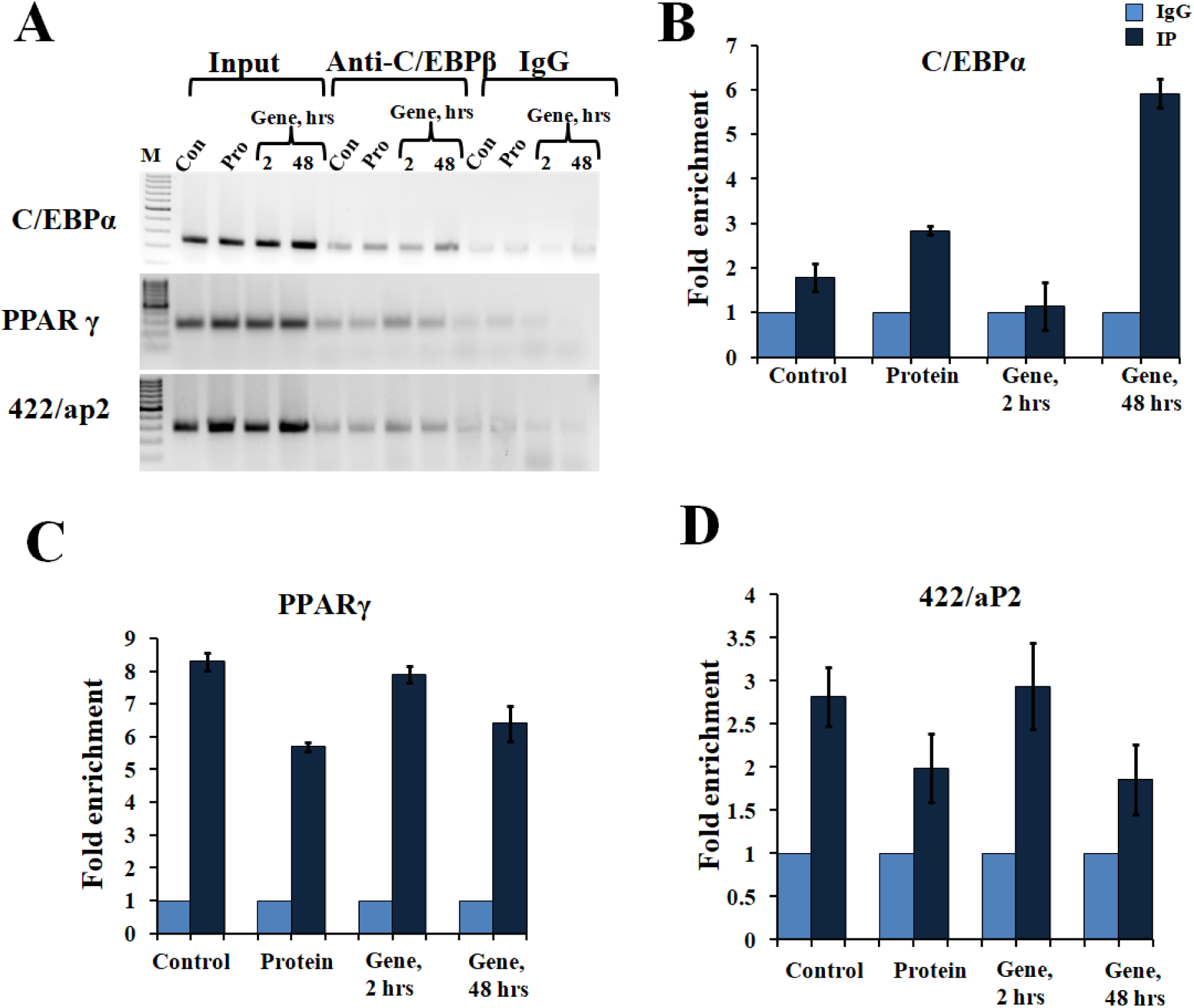
A-C/EBP transfection affects the binding of C/EBPβ to promoter region of adipocyte-marker genes. (**A**) Semi-quantitative PCR from ChIP samples showed binding of C/EBPβ to the C/EBP regulatory element in the promoter region of *C/EBPα, PPARγ, 422/aP2* in transfected cells. (**B-D**) Quantitative Real time PCR from ChIP samples of transfected cells with A-C/EBP protein and gene. Values are expressed as mean ±SD; n=3 independent experiments.

## 4 Discussion

Adipogenesis involving conversion of fat-free preadipocytes to lipid-laden adipocytes is a time-dependent differentiation phenomena that draws in many TFs during the process. This study was initiated with the assumption that the timely intervention and targeting TFs may inhibit preadipocytes differentiation, change cell fate, and reverse the process of differentiation. C/EBP family of bZIP TFs along with PPARγ are considered to be master regulators of adipocytes differentiation. We focused on inhibiting DNA binding activity and interactions of C/EBPβ with other factors by employing heterodimerzing protein inhibitor A-C/EBP. To achieve the objectives, A-C/EBP pure protein was directly delivered into 3T3-L1 cells using liposome delivery method. Parallelly, plasmid containing A-C/EBP gene was used as control. A-C/EBP was designed to inhibit the DNA binding of all the C/EBP family members [12]. Transgenic mice overexpressing A-C/EBP protein showed reduced level of C/EBPβ but elevated C/EBPα expression suggesting high *in vivo* specificity [41]. Here we show that A-C/EBP protein and gene transfected cells displayed reduced C/EBPβ mRNA and protein levels. C/EBPα mRNA level was down regulated with no change in protein expression whereas C/EBPδ levels remained impervious to A-C/EBP treatment. Liposome-mediated transfections successfully delivered A-C/EBP, an anionic peptide into 3T3-L1 cells as shown here and elsewhere by microscopy and biochemical approaches (Figure 1A in this study, [12]. Also, intense protein florescent signals were observed after 48hrs of plasmid transfection in the cells (Figure S2A).

Earlier, on-target efficiency of CRISPR/Cas9 nuclease system was evaluated by using pure Cas9 protein delivered directly into the cells and its activity was compared with mRNA and gene transfections where the presence of Cas9 protein was observed within hours of transfection whereas it took 24 hours to express the protein from DNA [43]. Similarly, in this study A-C/EBP protein signal peaked within 2hrs of transfection whereas it took 48hrs for A-C/EBP plasmids to express equivalent amount of protein (Figure 2A-D). A major advantage of using A-C/EBP is the formation of highly stable C/EBPβ|A-C/EBP heterodimer that is two magnitude stronger than C/EBPβ-DNA complex thereby precluding the requirement of A-C/EBP overexpression [12]. Furthermore, A-C/EBP specifically impacts the adipogenic pathway as 3T3-L1 cells transfected with A-VBP, an inhibitor of VBP bZIP TFs did not influence the differentiation process. Using A-C/EBP coding adenovirus, previous study showed the prevention of translocation of C/EBPβ into the nucleus [11]. Along similar lines, expression level of C/EBPβ from A-C/EBP protein transfected samples showed lower protein level after 2hrs, whereas A-C/EBP plasmid samples showed reduced C/EBPβ signals only after 48hrs of initiation of differentiation (Fig 5A-B). Degradation of C/EBPβ in early stages of differentiation led to lower expression of *PPARγ, C/EBPα*, 422/aP2 and other adipocyte-marker genes.

We further analyse the expression profiles of known C/EBP interacting partners. Stat5a is known to co-localize with C/EBPβ whereas CREB binds to the promoter region of *C/EBPβ* and GATA2/3 form protein complexes with C/EBPβ/α [30–32]. Also, STAT5a and CREB co-express at the same time as C/EBPβ i.e., after an hour of induction whereas GATA2/3 are expressed at the onset of adipogenesis [31,32,44]. With the degradation of C/EBPβ at early stage of differentiation there is a concomitant decrease in the expression of its direct interacting partners as indicated by lower expression of *Stat5a* and *CREB*. Observed higher *GATA2* expression in protein transfected cells is may be due to the sequestering of C/EBPβ by A-C/EBP. GATA2 is known to bind to C/EBPβ/α In absence of C/EBPβ it binds to C/EBPα leading to suppression of adipogenesis in protein transfected cells. In A-C/EBP plasmids sample no change in expression was observed for GATA 2/3 whereas in 48hrs GATA3 mRNA was enhanced without any change in GATA 2 level. This observation reinforces the advantages of using proteins and peptides to study the spatial and temporal gene regulation in complex pathways like adipogenesis. *TCF7L1* and *KLF2* are not known to interact with C/EBPβ and their upregulation in A-C/EBP gene but not in protein transfected samples may be due to non-specific interactions of A-C/EBP at higher and persistent expression.

We went on to explore if we could revert cell fate and de-differentiate adipose cells. Figure 7 shows the experimental design and summary results obtained when 3T3-L1 cells were transfected with A-C/EBP protein and gene. De-differentiation of mature adipocyte during lactation into preadipocyte precursor cells have shown expression of common preadipocyte-markers i.e., *Pdgfrβ, Pdgfrα, Ly6A* (*Sca1*), *CD34* and *Itgb1*(fibronectin receptor) [39]. Gene expression analysis of common preadipocyte-markers from cells post 5-days of differentiation, have shown higher expression of all five genes in A-C/EBP protein transfected cells. In contrast, we observed the non-significant expression of *Pdgfrβ, Pdgfrα* and *Itgb1* but higher expression of *Ly6A* and *CD34* in gene (48hr) transfected cells. *Ly6A, CD34* and *Itgb1* are the common mesenchymal stem cell markers but the signalling balance between *Pdgfrβ/α* regulate the commitment of adipose progenitors to either white or beige formation [38,40]. Earlier studies have shown the inhibitory role of *Pdgfrα* during adipogenesis as its activation leads to inhibition of adipogenesis [45,46]. Also mosaic deletion of both *Pdgfrα* and *Pdgfrβ* enhances the adipogenesis process as these act as negative regulator of adipogenesis [47]. Downregulation and degradation of Pdgfrs within 6hrs of hormonal induction has been observed in 3T3-L1 cells [48]. C/EBPβ is reported to transcriptionally repress *Pdgfrα* in vascular smooth muscle cells (VSMCs) [49]. Akin to VSMCs, our results show that C/EBPβ inhibition by A-C/EBP protein upregulate the expression of *Pdgfrα/β* after hormonal induction thus inhibiting the conversion of preadipocytes to adipocytes. Our findings suggest that higher expression of common preadipocyte-markers may be due to non-conversion of cells into adipocytes even after hormonal induction. In gene transfection studies, late expression of A-C/EBP protein though inhibits the differentiation process but did not induce all preadipocytes marker genes. A-C/EBP gene transfected cells show similar morphology as that of protein transfected cells since late expression of A-C/EBP leads to suppression of C/EBPβ which further inhibit the expression of C/EBPα and PPARγ, two master regulators of terminal adipocyte formation. Thus, the strategy to introduce exogenous protein directly to the mammalian cells may control the fate of differentiating cells and may be applied to study signal transduction that involved co-ordinated and timely expression of number of TFs. Elaborate and quantitative proteomic studies will decipher the absolute and relative expression levels of preadipocytes markers that decide the cell fate, for example, relative expression of Pdgfrα/Pdgfrβ defines the cell fate with lower ratio leading to WAT (white adipose tissue) and higher ratio to BAT (beige adipose tissue).

**Figure 7.**
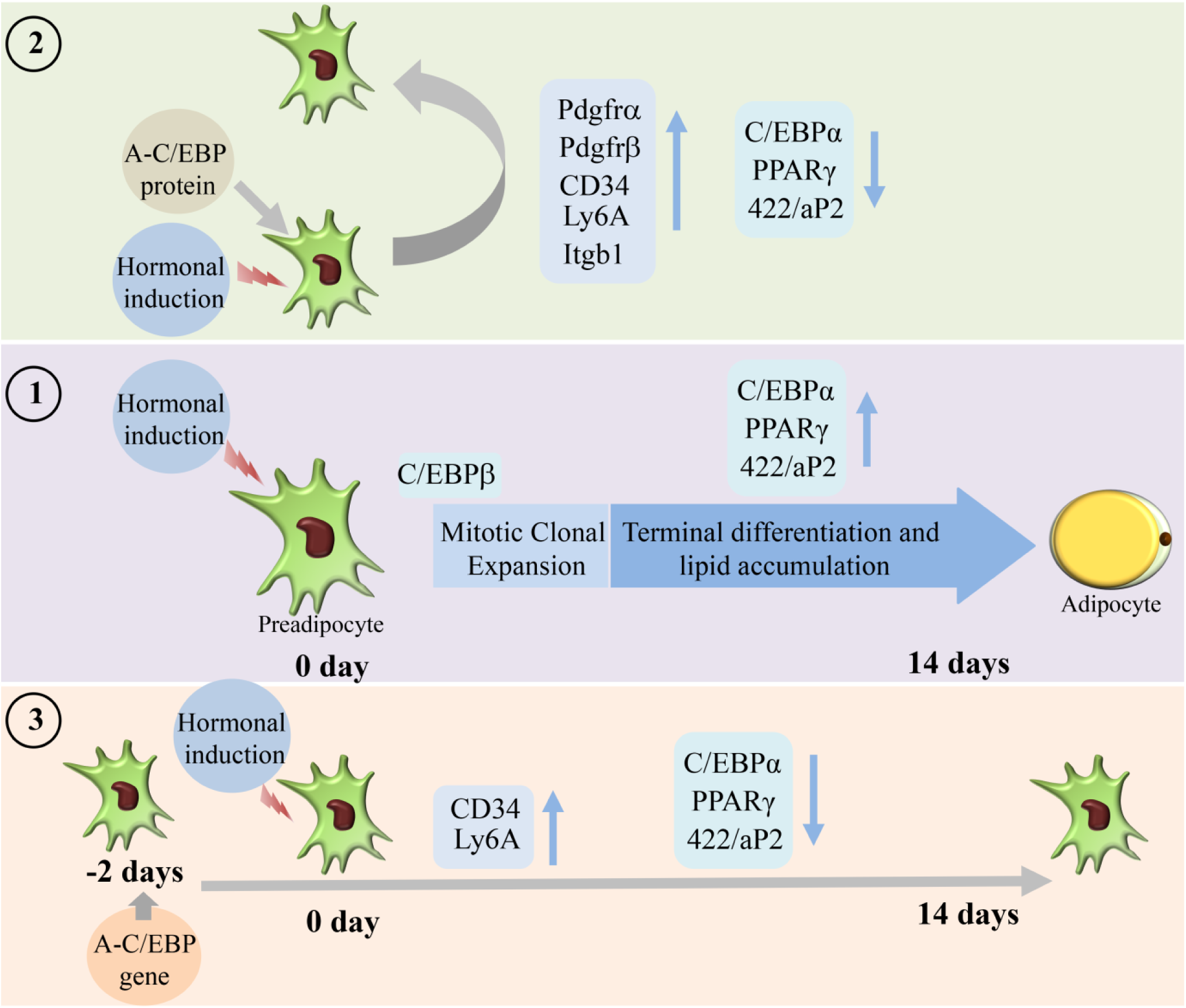
Summary of the studies showcasing the results obtained when 3T3-L1 cells were transfected with A-C/EBP protein and gene. Three experimental conditions are depicted here (**1**) Normal differentiation protocol was followed. 0 day defines the day when cells were exposed to MDI cocktail of hormones. Differentiation process takes 14 days that involves mitotic clonal expansion and terminal differentiation. Process is marked by transformation of flaccid preadipocytes into lipid-laden mature adipocytes. Only master regulator of adipogenesis e.g., C/EBPβ, C/EBPα, PPARγ, and 422/aP2 are shown. (**2**) Cells were transfected with A-C/EBP protein for 2-6 hrs and were induced to differentiate. Following differentiation protocol, cells were tested for phenotype and analysed for molecular markers. Cells failed to differentiate as determined by Oil Red O staining. Cells transfected with A-C/EBP protein still express preadipocyte markers 5-days post differentiation. Five preadipocyte markers were upregulated, among them Pdgfrα and Pdgfrβ are responsible for white adipocyte tissue (WAT) to beige adipocyte cell fate determination. (**3**) Cells were transfected with A-C/EBP gene for 48hrs and differentiated. After 14 days cells were stained with Oil Red O and adipocyte differentiation markers were analysed. Unlike protein samples only two CD34 and Ly6A preadipocytes markers were expressed.

## Acknowledgements

We thank Executive Director, National Agri-Food Biotechnology Institute (NABI), Mohali for research facilities and Department of Biotechnology (DBT), New-Delhi for research funding. NS acknowledges the financial assistance from NABI and DBT in the form of JRF and SRF.

## Author Contributions

N.S., and V.R. designed the study. N.S., and V.R. analyzed the data and wrote the manuscript. N.S. conducted the experiments with help from R.K., K.S., H.P., D.C., P.J., and A.A. Confocal microscopy experiments were performed with help from B.Y. C.V. provided the reagents. All authors reviewed the results and approved the final version of the manuscript.

## Legend to Supplementary Tables

**Table S1**: Primers for qRT-PCR of adipocyte -marker genes and genes involved in adipogenesis.

**Table S2**: Primers for semi-quantitative ChIP-PCR and ChIP-qPCR.

**Table S3**: Primers for preadipocyte-marker genes.

