## Supplementary figures and images for "Transient delivery of A-C/EBP protein perturbs differentiation of 3T3-L1 cells and induces preadipocyte marker genes"

### Supplementary figures

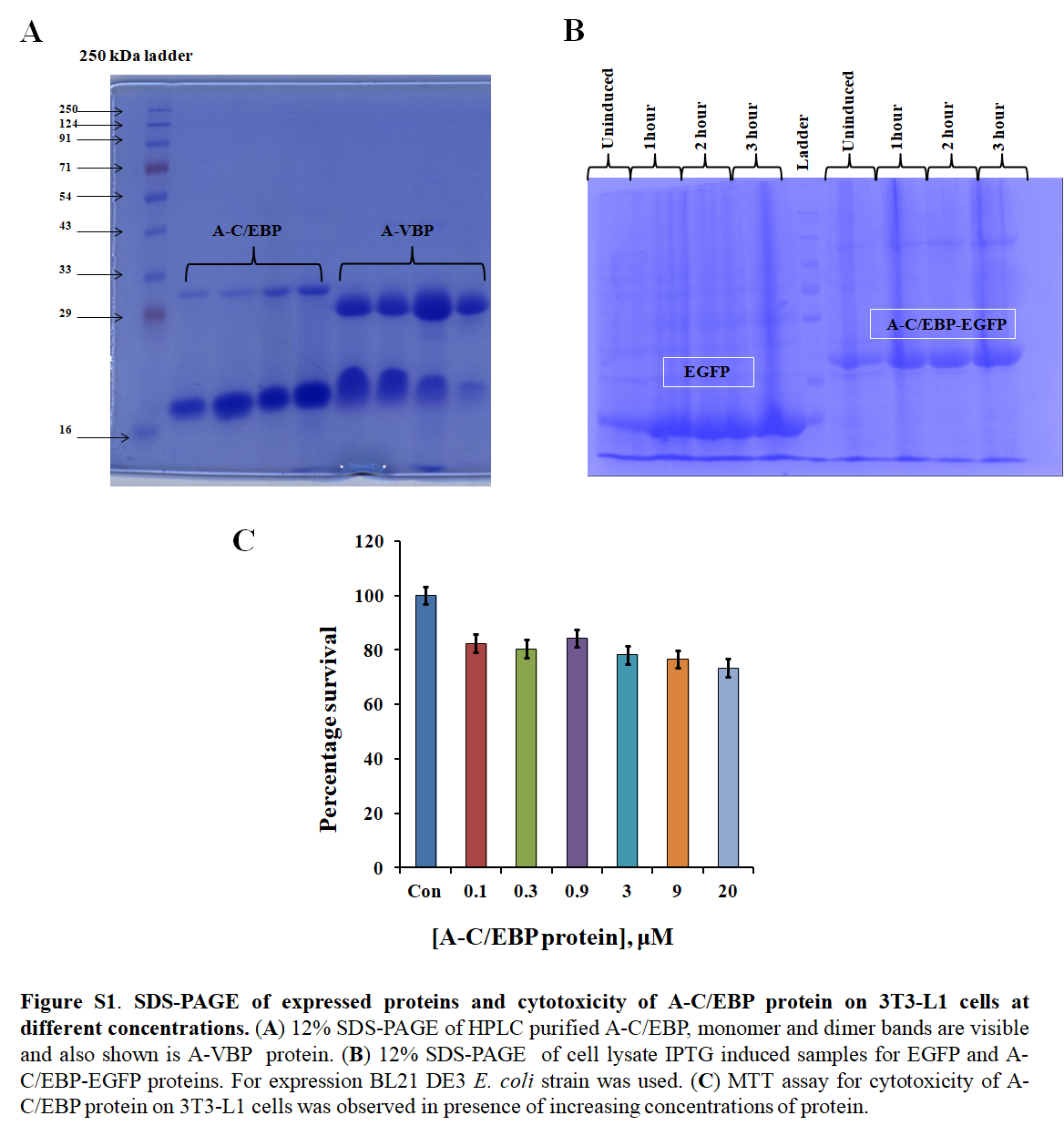

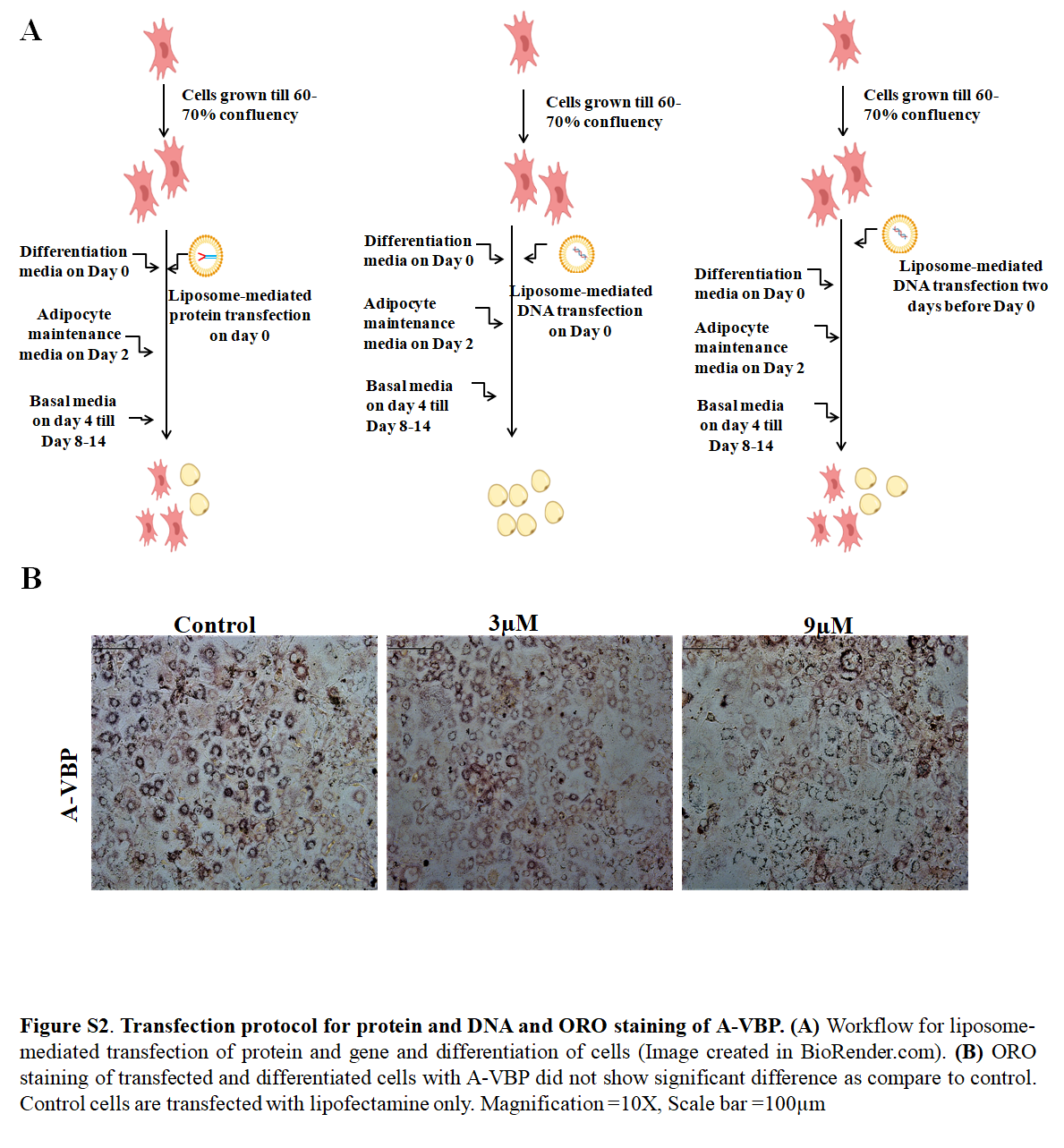

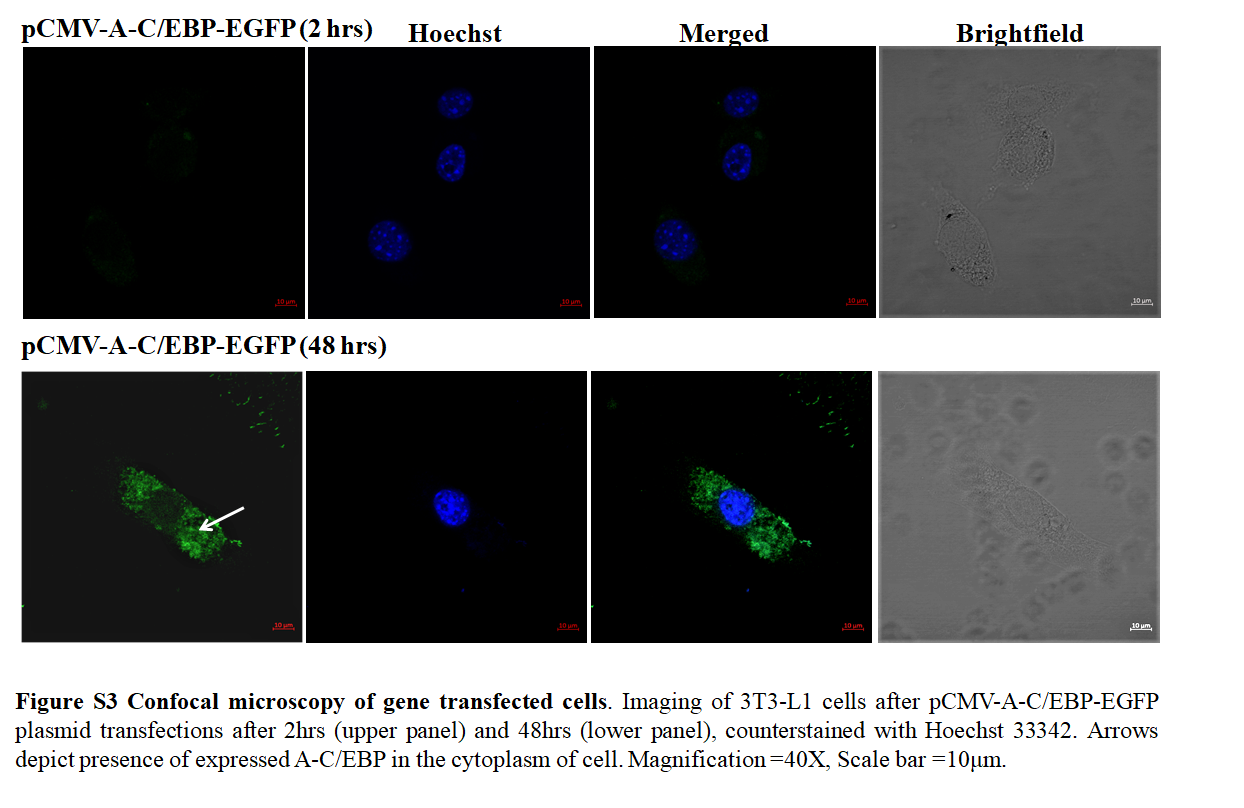
