## Supplemental Tables for "Transient delivery of A-C/EBP protein perturbs differentiation of 3T3-L1 cells and induces preadipocyte marker genes"

**Supplementary**

**Table 1. Primers for qRT PCR of adipocyte -marker genes and genes involved in adipogenesis**

| **Gene** | **Gene ID** | **Forward primer** | **Reverse primer** | **Amplicon size(bp)** |
| --- | --- | --- | --- | --- |
| β-actin | NM_007393 | CTCTGGCTCCTAGCACCATGAAGA | GTAAAACGCAGCTCAGTAACAGTCCG | 200 |
| C/EBP β | NM_009883.4 | AGCGACGAGTACAAGATGCG | TCAGCTCCAGCACCTTGT | 109 |
| C/EBPδ | X61800.1 | GCAACAACATCGCTGTGC | TCCACGCGCTGATGCAGCTT | 115 |
| C/EBPα | NM_001287523.1 | ATCGATCCATCCCAGAGGGA | ATAGACGTGCACACTGCCA | 132 |
| PPARγ | NM_001127330.2 | TGGTGTCCATGAGATCATCT | GGAACTCCCTGGTCATGA | 98 |
| Fabp4 | NM_024406.3 | TGAAATCACCGCAGACGACAGG | CACCAGCTTGTCACCATC TCG | 131 |
| Scd1 | NM_009127.4 | TTGGTGGGAAGTAATTCGAG | TTCAGAGACTACACCCAGAG | 117 |
| GLUT4 | NM_011400.3 | AGCTAGATGAGACCTCTTCC | TTCTGGAGCCATCAAAGTC | 113 |
| SREBP1 | AF374266.1 | AGTCACCAGCTTCAGTCCAG | CTCAGGTCATTTTGGAAACC | 112 |
| KLF6 | NM_011803.2 | GTTTCTGCTCGGACTCCTGAT | TTCCTGGAAGATGCTACACATTG | 108 |
| Stat5a | Z48538.1 | CAGCCGTGGGATGCTATTGA | GGGACAGCGGTCATACGTG | 186 |
| Zfp423 | NM_033327.2 | TGGCCTGGGATTCCTCTGT | CTCTTGACTTGTCACGCTGTT | 110 |
| Zfp467 | NM_001085416.1 | TCCTGCTCAGGGCATGAGA | TCCGAATCATCCATTCCTCCC | 145 |
| TCF7L1 | NM_001079822.2 | ACGAGCTGATCCCCTTCCA | CAGGGACGACTTGACCTCAT | 101 |
| CREB | U46027.1 | CTGAGAGCTGGTATGTCAGGA | TGAGTGCTGGAGTAAAACAGTCA | 125 |
| Pref1 | NM_010052.5 | TTCGGCCACAGCACCTATG | GGGGCAGTTACACACTTGTCA | 134 |
| KLF2 | NM_008452.2 | GAGCCTATCTTGCCGTCCTTT | CACGTTGTTTAGGTCCTCATCC | 117 |
| GATA2 | NM_008090.5 | CACCCCGCCGTATTGAATG | CCTGCGAGTCGAGATGGTTG | 130 |
| GATA3 | NM_008091.3 | CCCCATTACCACCTATCCGC | CCTCGACTTACATCCGAACCC | 106 |
| FoxC2 | NM_013519.2 | ACGTCCGGGAGATGTTCAAC | GACGGCGTAGCTCGATAGG | 109 |
| VEGF | AY707864.1 | GCACATAGAGAGAATGAGCTTCC | CTCCGCTCTGAACAAGGCT | 105 |
| Cyclin A | Z26580.1 | GCCTTCACCATTCATGTGGAT | TTGCTGCGGGTAAAGAGACAG | 118 |
| Cdk2 | NM_016756.4 | ATGGAGAACTTCCAAAAGGTGG | CAGTCTCAGTGTCGAGCCG | 124 |
| Cyclin B1 | NM_172301.3 | GCGTGTGCCTGTGACAGTTA | CCTAGCGTTTTTGCTTCCCTT | 135 |
| Cyclin D1 | NM_007631.2 | GCGTACCCTGACACCAATCTC | ACTTGAAGTAAGATACGGAGGGC | 94 |
| p27 | U10440.1 | TCAAACGTGAGAGTGTCTAACG | CCGGGCCGAAGAGATTTCTG | 103 |

**Table 2. Primers for semi-quantitative ChIP-PCR and ChIP-qPCR**

| **S.no**. | **C/EBP binding site in:** | **Forward primer** | **Reverse primer** |
| --- | --- | --- | --- |
| **1.** | C/EBPα promoter | TGGAAGTGGGTGACTTAGAGGC | GGATGGTGCCTGCTGGGTCTTA |
| **2.** | PPARγ promoter | TTCAGATGTGTGATTAGGAG | AGACTTGGT ACATTACAAGG |
| **3.** | 422/aP2 promoter | CCTCCACAATGAGGCAAATC | CTGAAGTCCAGATAGCTC |

**Table 3. Primers for preadipocyte-marker genes**

| **Gene** | **Gene ID** | **Forward primer** | **Reverse primer** | **Amplicon size** |
| --- | --- | --- | --- | --- |
| Pdgfrb | NM_008809.2 | AGGAGTGATACCAGCTTTAGTCC | CCGAGCAGGTCAGAACAAAGG | 152 |
| Pdgfra | NM_001083316.2 | ATGAGAGTGAGATCGAAGGCA | CGGCAAGGTATGATGGCAGAG | 130 |
| Ly6A (Sac1) | NM_001271416.1 | GCTCAGTCCTCCTGCAGACCT | TTCACACACTACTCCCACCTT | 146 |
| CD34 | NM_133654.3 | CTGGGTAGCTCTCTGCCTGAT | TGGTAGGAACTGATGGGGATATT | 101 |
| Itgb1 (CD29) | NM_010578.2 | ACTGTGATGCCGTATATTAGCAC | GATATGCGTTGCTGACCAACA | 155 |
